# Genetic control of rhizosheath formation in pearl millet

**DOI:** 10.1101/2021.11.02.466908

**Authors:** C. De la Fuente Cantó, M.N. Diouf, P.M.S. Ndour, M. Debieu, A. Grondin, S. Passot, A. Champion, C. Barrachina, M. Pratlong, P. Gantet, K. Assigbetsé, N. Kane, P. Cubry, A.G. Diedhiou, T. Heulin, W. Achouak, Y. Vigouroux, L. Cournac, L. Laplaze

## Abstract

The rhizosheath, the layer of soil that adheres strongly to roots, influences water and nutrients acquisition. Pearl millet is a cereal crop that plays a major role for food security in arid regions of sub Saharan Africa and India. We previously showed that root-adhering soil mass is a heritable trait in pearl millet and that it correlates with changes in rhizosphere microbiota structure and functions. Here, we studied the correlation between root-adhering soil mass and root hair development, root architecture, and symbiosis with arbuscular mycorrhizal fungi and we analysed the genetic control of this trait using genome wide association (GWAS) combined with bulk segregant analysis and gene expression studies. Root-adhering soil mass was weakly correlated only to root hairs traits in pearl millet. Twelve QTLs for rhizosheath formation were identified by GWAS. Bulk segregant analysis on a biparental population validated five of these QTLs. Combining genetics with a comparison of global gene expression in the root tip of contrasted inbred lines revealed candidate genes that might control rhizosheath formation in pearl millet. Our study indicates that rhizosheath formation is under complex genetic control in pearl millet and suggests that it is mainly regulated by root exudation.

**Highlight:** Formation of the rhizosheath, a layer of soil adhering to the root, is under complex genetic control in pearl millet and is mainly regulated by root exudation.

## Introduction

Pearl millet is a small-seeded tropical cereal that was domesticated about 4,500 years ago in the Sahelian part of West Africa (Burgarella *et al*., 2018). It is mostly grown in dry and poor soils as a rainfed crop and is therefore well adapted to environments prone to drought and heat stress for which it harbours largely untapped genetic diversity in the locally adapted cultivated and wild pearl millets (Debieu *et al*., 2017; Varshney *et al*., 2017; Burgarella *et al*., 2018). The outstanding capacity for growing in harsh environments highlights the great potential of pearl millet as a biological model to investigate crop adaptation and resilience to abiotic constraints, as well as its key role for food security in some semi-arid tropical regions in Africa and Asia. Still, pearl millet yield remains low for two main reasons: the difficulty to reach its potential yield in constrained environments and the little attention that the crop has received from breeding programmes (Varshney *et al*., 2017).

Root traits are emerging as new targets for breeding more sustainable and resilient crop varieties in global climate change scenarios (Lynch, 2019). The root system is responsible for plant water and nutrient acquisition. Phenotypic selection of root ideotypes combining architectural, anatomical and physiological traits has been proposed as a way to optimise access to soil resources in specific agroecosystems and crop management practices (Lynch, 2019). Besides root architecture, anatomy and physiology, the rhizosphere, the volume of soil around the root influenced by the root (York *et al*., 2016), can be regarded as a plant extended phenotype and therefore a target for breeding more sustainable crops (Wissuwa *et al*., 2009; De la Fuente Cantó *et al*., 2020). Indeed, the dynamic interplay between root, soil and microbiota in the rhizosphere eases adaptation to changing environments and can have a remarkable impact on plant fitness (Turner *et al*., 2013; De la Fuente Cantó *et al*., 2020; Chai and Schachtman, 2021). The intricate relationships in the rhizosphere define a belowground niche where soil moisture, organic matter content, the composition of the microbial community and its activity are different from the bulk soil (Haichar *et al*., 2008; Hinsinger *et al*., 2009). Plants benefit from this interaction especially in constrained environments where access to nutrients and water is restricted (Yang *et al*., 2009).

The rhizosheath size, or root-adhering soil mass, is a proxy in the study of this complex extended phenotype and an interesting potential target for breeding programmes (Ndour et al., 2020). Rhizosheath formation was first noticed as the sandy sheath surrounding the roots of desert plants (Price, 1911) and later reported to occur across many angiosperm orders (Brown *et al*., 2017). Increased rhizosheath size has been correlated with enhanced wheat and foxtail millet performance in drying soils (Basirat *et al*., 2019; Liu *et al*., 2019). In barley and oat, rhizosheath formation has been related with improved acquisition of major and essential trace elements in limiting water conditions (Nambiar, 1976; George *et al*., 2014). A combination of root architectural and anatomical traits and the secretion of root exudates and mucilage have been connected to soil aggregation to the root (Pang *et al*., 2017; Ndour *et al*., 2020). For instance, root branching, root hair formation or symbiosis with arbuscular mycorrhizal fungi (AMF) have been associated to some extent with rhizosheath establishment (Moreno-Espíndola *et al*., 2007; Brown *et al*., 2017; Liu *et al*., 2019). Root architectural traits have been found crucial for rhizosheath formation in wheat and foxtail millet (Delhaize *et al*., 2012; Liu *et al*., 2019). On the other hand, root exudates composition and mucilaginous polymers released by root-associated microorganisms impact the stability of soil aggregates that bind around the root (Galloway *et al*., 2020). Root growth and exudates exert dynamic changes in the rhizosphere physical properties and hydraulic processes that affect soil nutrient dynamics and the composition of the rhizosphere associated microbiota (Dakora and Phillips, 2002; Kolb *et al*., 2017; Sasse *et al*., 2018; Chai and Schachtman, 2021). Despite the inherent complexity linked to the effect of exudates in the rhizosphere, some studies showed their direct relationship with rhizosheath formation. For example, greater mass of mucilage exuded by chickpea roots were linked with the formation of larger and more porous rhizosheaths capable of storing more soil moisture in drought tolerant cultivars (Rabbi *et al*., 2018). In annual crops such as wheat, barley and maize, there is evidence of the remarkable plant genetic influence in the formation of rhizosheath and the processes of rhizodeposition influencing rhizosphere microbial activities (George *et al*., 2014; Delhaize *et al*., 2015; Mwafulirwa *et al*., 2016; 2021b), however few studies have aimed to dissect the genetics underlying the conformation of this extended root phenotype (George *et al*., 2014; Delhaize *et al*., 2015; James *et al*., 2016; Mwafulirwa *et al*., 2021a).

In previous studies, we reported a remarkable genotypic variability for root-adhering soil aggregation in pearl millet (Ndour *et al*., 2021). Moreover, this variability was associated with changes in rhizosphere microbiota structure and function (Ndour *et al*., 2017, 2021). Here, we analysed the relative contribution of root architectural characteristics and root colonization by AMF on root-adhering soil aggregation in pearl millet. We then combined a genome wide association analysis (GWAS), with bulk segregant analysis (BSA) and transcriptomic data to dissect the genetic bases of this complex trait.

## Materials and methods

### Plant materials

A panel of 181 pearl millet inbred lines developed at the International Crops Research Institute for the Semi-Arid Tropics (ICRISAT, Niger) from landraces and improved open-pollinated cultivars representing the genetic diversity of the crop in West and Central Africa was used in this study (Debieu *et al*., 2018).

Two inbred lines from this panel with contrasted rhizosheath size measured by the ratio between the mass of root-adhering soil (RAS) and root biomass (RT; RAS/RT ratio; Ndour *et al*., 2021): ICML-IS 11139 (small rhizosheath size parent) and ICML-IS 11084 (large rhizosheath size parent) were selected for a bi-parental cross. The obtained F2 offspring was then used in a bulk segregant analysis (BSA).

### Plant growth and measurement of soil aggregation

Plants were grown for 28 days in “WM” shaped pots (WM 20-8-5, Thermoflan, Molières-Cavaillac) containing 1.5 kg of soil under natural light in a greenhouse in the ISRA/IRD Bel Air Campus in Dakar (Lat. 14.701778, Long. −17.426229, altitude 9 m) as previously described (Ndour *et al*., 2021).

For the GWAS analysis, pearl millet lines were sown according to a complete random block design with seven repetitions. Thinning was performed to have one plant per pot. Soil moisture was adjusted daily at water-holding capacity. Plant watering was stopped 24 hours before harvesting to facilitate the separation of root-adhering soil (RAS) from bulk soil. Plants were harvested 28 days after planting by opening the pots gently and shaking the plant and its adhered soil at a constant speed (1100 rpm) for 1 min with a CAT S50 electric shaker (Cat Ingenieurbuero™) to separate the bulk soil from the RAS uniformly. Roots were then rinsed in a cup with demineralized water to collect RAS. The RAS was dried at 105 °C for three days and weighted. Roots and shoots were separated and dried at 65 °C for three days. The ratio between mass of RAS and mass of root tissue (RT; RAS/RT) was used to estimate the rhizosphere aggregation intensity (rhizosheath size) as previously described (Ndour *et al*., 2017).

For BSA analysis, RAS/RT ratio was measured on 553 F2 individuals grown in five successive blocks of 119, 112, 131, 130 and 61 F2 plants. Each of these blocks included six replicates randomly distributed for each parental line. At the end of the experiment, leaf disk samples of 1.5 mm diameter were sampled for each individual plant and stored at −80 °C for genotyping.

For correlation analyses between RAS/RT ratio and related root traits (root architecture, root hair length and density and interactions with AMF), 8 contrasting genotypes for rhizosheath size were analysed in 2018 and 2020 (n=10 plants/genotype in 2018 and n=6 plants/genotype in 2020).

### Root architecture

Root architecture traits (length, average diameter, total root area) were measured using the WinRHIZO software (version 2012b) after scanning the roots using an Epson Perfection V700 scanner. Roots were separated in two groups based on their diameter according to Passot *et al*. (2016): primary and crown roots (0.25 mm < diameters) and lateral roots (diameters < 0.25 mm).

Root hair length and density were measured on four plants per genotype using images of the root hair zone of three lateral roots per plant. Images were taken using an optical microscope (BX50F, Olympus) equipped with a digital camera (Micro Publisher 3.3 RTV). For each lateral root, the total number of root hairs was recorded over a distance of 0.5 mm using the Mesurim free software (http://acces.ens-lyon.fr/acces/logiciels/applications/mesurim/mesurim) and the length of 10 randomly selected root hairs was measured using the ImageJ software.

### Root colonization by arbuscular mycorrhizal fungi

Intensity and frequency of root colonization by AMF were measured according to Trouvelot *et al*., (1986) after roots staining with Trypan blue following the method described by Phillips and Hayman (1970). Stained root fragments were observed with a Nikon Labophot trinocular microscope. For each fragment, a score between zero and five was assigned according to the estimated proportion of root cortex colonized by AMF (Trouvelot *et al*., 1986).

Frequency and intensity of root colonization were then computed using the following formulas:

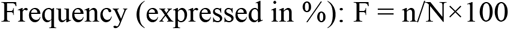

where n is the number of fragments showing mycorrhizae and N, the number of observed fragments

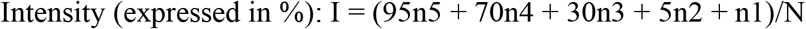

where n1, n2, n3, n4, n5 are the number of fragments scored respectively from 1 to 5 and N, the number of fragments observed.

### Heritability

Broad sense heritability was computed using the following formula:

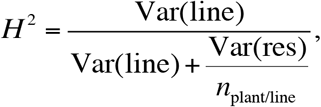

where *η*_plant/line_ is the average number of plants measured per line, Var(line) is the variance associated with lines and Var(res) is the residual variance.

Both variances are parameters of the following linear mixed model:

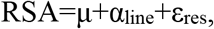

where *μ* is the overall mean soil aggregation, *α*_line_ is the random effect attached to the lines with *α*_line_ ~ N(0, Var(line)) and *ε*_res_ is the error term with *ε*_res_ ~ N(0, Var(res)).

### Genome wide association mapping and statistical analysis

Genotyping by sequencing of this panel of inbred lines was previously reported (Debieu *et al*., 2018). As a preliminary step, we used the genotypic matrix to estimate the population structure. Individual ancestry coefficients were estimated using the R package LEA v2.0 (Frichot and François, 2015). We used a latent factor mixed model (LFMM) that considers ridge estimates and corrects for unobserved population cofounders, *i.e*. latent factors, to perform the GWAS (Caye *et al*., 2019). In addition, we ran the efficient mixed-model association (EMMA, Kang *et al*., 2008) and mixed linear model (MLM) implemented in the R package GAPIT (Lipka *et al*., 2012) to contrast the results. The proportion of the phenotypic variance explained by a QTL was determined by estimating the R^2^ corrected for population structure of a linear model defined for the most significant SNPs.

### Bulk Segregant Analysis

NGS-based BSA studies require establishing contrasted groups or bulks of lines to assess the differences in segregation of alleles using high-throughput sequencing (K. L. Nguyen *et al*., 2019). The 10% extreme lines in the tails of the phenotype distribution for the RAS/RT ratio were selected and the corresponding leaf discs were pooled to form bulks of contrasted lines. Genomic DNA was isolated for each bulk using a MATAB (Mixed Alkyl Trimethyl Amonium Bromide) based method (Mariac *et al*., 2006) and enriched DNA libraries were constructed for which 32,860 predicted genes from the pearl millet reference genome (Varshney *et al*., 2017) were targeted using gene capture probes (myBaits^®^). High-throughput sequencing of the enriched DNA library was performed on an Illumina HiSeq platform by Novogene Company Limited (HK). Initial sequencing quality checks using FastQC version 0.11.5 (Andrews, 2010) were followed by trimming and quality filter steps on which adaptors, barcode sequences and low-quality reads (< 35 bp) were removed. Paired sequences were then retained and aligned to the pearl millet reference genome using the BWA MEM algorithm (BWA version 0.7.17 - r1188, Li and Durbin, 2009). Reads mapping at the target-enriched regions were used for SNP calling using the UnifiedGenotyper algorithm from GATK 3.7 (McKenna *et al*., 2010). Down-sampling limit (dcov) was increased from the default value of 250 to 9000 to ensure accounting for the maximum coverage reached at each position. Multi-allelic sites and those which exhibited a total allele frequency less than 0.25 were removed. In addition, sites with either low or high total sequencing depth (below the 25^th^ and above the 95^th^ percentiles respectively) were removed. SNPs with more than 50% missing data and minor allele frequency (MAF) under 5% were also excluded. Finally, the parental line ICML-IS 11139 (low RAS/RT ratio) was used as the reference genome for the cross to designate the alternate and reference SNP variants in the bulks.

Euclidean distance-based statistics (Hill *et al*., 2013) was used to measure the difference in allele frequency between the bulks. The Euclidean distance between allele frequencies of the bulks at each marker position (*EDm*) was calculated as follows:

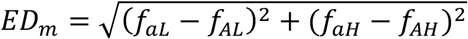

where *f_a_* and *f_A_* correspond to the allele frequency of the alternate and reference allele in the low bulk (L) and the high bulk (H) respectively.

In order to reduce the effect of sequencing noise and increase the signal of the differences in allele frequency, we then calculated the fourth power of the cumulative *EDm* value in windows of 100 consecutive markers (Omboki *et al*., 2018; Zhang *et al*., 2019). The confidence interval of the statistic was determined using simulations as described (de la Fuente Cantó and Vigouroux 2021, under revision)

### Gene expression analyses

Seeds from lines ICML-IS 11139 (low RAS/RT ratio) and ICML-IS 11155 (high RAS/RT ratio) were surface-sterilized and germinated in Petri dishes containing wet filter paper for 24 h in the dark at 27 °C. After two days, plants were transferred to hydroponic tanks containing liquid half Hoagland solution and grown for 15 days at 27 °C (12 h light/12 h dark). RNA was extracted from crown root tips (two cm apex) using the RNeasy Plant Mini Kit (QIAGEN). RNA-seq was performed by the Montpellier GenomiX Platform (MGX, https://www.mgx.cnrs.fr/). Sequencing was performed on an Illumina HiSeq 2500. Three different statistical tests were used to identify differentially expressed genes: EdgeR (Robinson et al., 2010), DESeq (Anders and Huber, 2010) and DESeq2 (Love *et al*., 2014). GO terms enrichment was performed in the 1270 genes that were significantly differentially expressed between the two lines for all three statistical tests using the TopGO package in R.

### Statistical methods

All statistical analyses were performed with R version 3.6.3 (R core Team, 2018).

## Results

### Root-adhering soil aggregation is weakly correlated to root hair traits in pearl millet

Several root traits have been proposed to contribute to root-adhering soil aggregation (as an integrative phenotype) including root hair development, root architecture, and arbuscular mycorrhizal symbiosis. We therefore analysed the contribution of these different traits to root-adhering soil aggregation in pearl millet. For this, we analysed correlation between root-adhering soil aggregation, root architecture, root hair length and density and frequency and intensity of root colonization by AMF in eight inbred lines with contrasted rhizosphere aggregation phenotype (Ndour *et al*., 2021) after four weeks of growth. Among root architecture traits, only the average root diameter (AvgDiam) was weakly and negatively correlated (*p* = 0.012, *r^2^* = 0.057) with root-adhering soil aggregation (Fig. 1A, Table 1). For root hairs, only average length of root hairs (AvgLRH) was weakly and positively correlated to root-adhering soil aggregation (*p* = 0.005, *r^2^* = 0.077; Fig. 1A, Table 1). No significant correlation was observed for all other traits including frequency and intensity of root colonization by AMF (Fig. 1A, Table 1). Similar results were found in two independent experiments (2018 and 2020; Supplementary Table S1 at *JXB* online).

**Figure 1.**
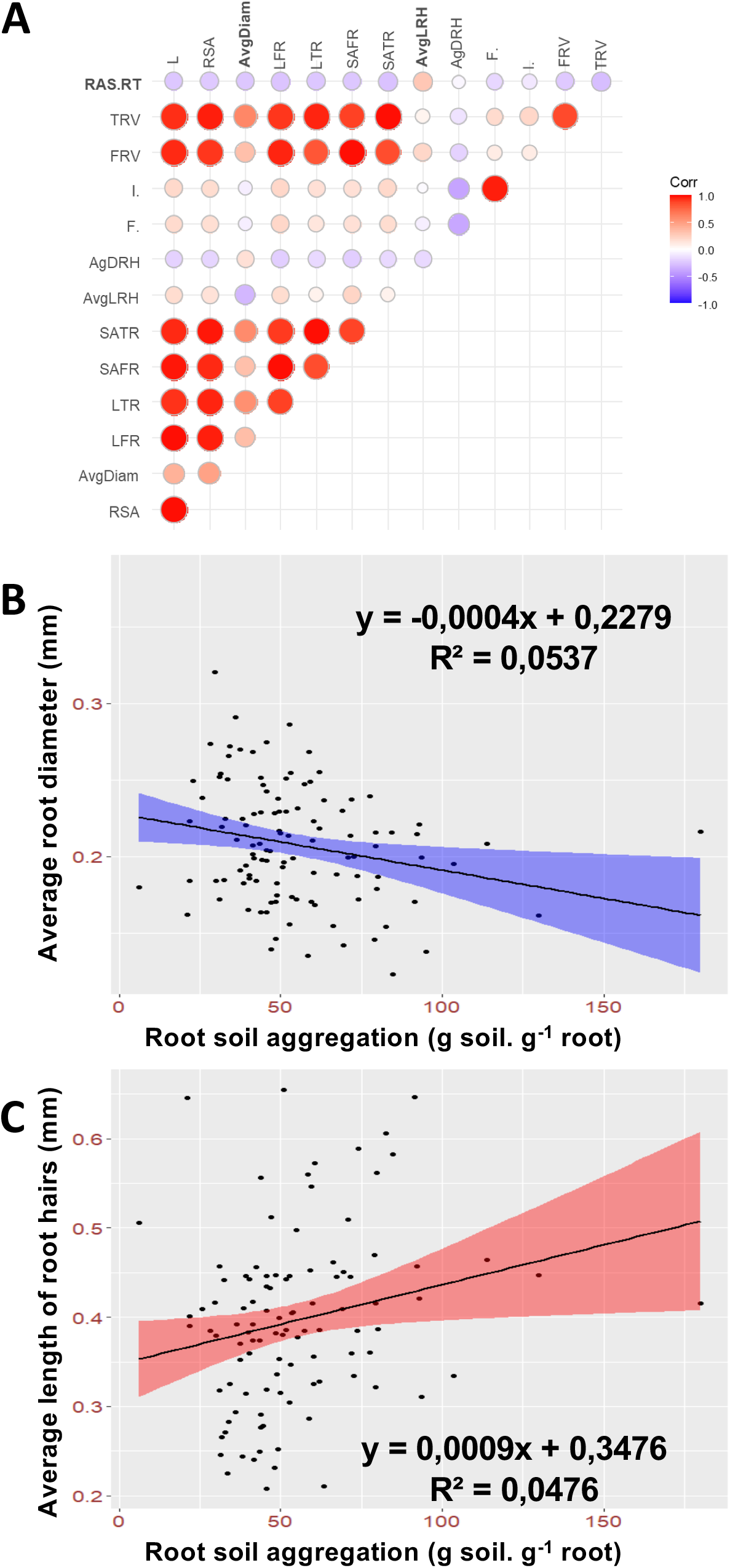
Relation between root soil aggregation, root architecture, root hair development and arbuscular mycorrhizal symbioses. **A)** Pearson correlation between traits using adjusted lsmeans across two experiments conducted in different years. **B)** Linear regression between root diameter and root soil aggregation. Points represent the mean value of the traits for inbred lines across the two experiments. **C)** Linear regression between root hair length and root soil aggregation. Points represent the mean value of the traits for inbred lines across the two experiments.

**Table 1.** Correlation matrix of root soil aggregation and root parameters.

| Trait | Root architecture |  |  |  |  |  |  |  | Root hairs |  | AM symbiosis |  |
| --- | --- | --- | --- | --- | --- | --- | --- | --- | --- | --- | --- | --- |
|  | RAS/R<br>T | L | RSA | AvgDiam | LFR | LTR | SAFR | SATR | AvgLRH | AgDRH | F% | I% |
| **RAS/RT** | **1** | 0,017 | -0,081 | **-0,239** | 0,053 | -0,112 | 0,037 | -0,110 | **0,278** | 0,069 | 0,283 | -0,133 |
| *p*-value | **0** | 0,863 | 0,397 | **0,012** | 0,579 | 0,242 | 0,697 | 0,250 | **0,005** | 0,488 | 0,463 | 0,744 |

Value from 2 independent experiments on contrasted pearl millet lines using Spearman's correlation test. Ratio (RAS/RT) between the mass of root-adhering soil (RAS) and root tissue biomass (RT), Total root length (L), Root Surface Area (RSA), Average Root Diameter (AvgDiam), Total Length of Fine Roots (LFR), Total Length of Thick Roots (LTR), Surface Area of Fine Roots (SAFR), Surface Area of Thick Roots (SATR), Average Length of Root Hairs (AvgLRH), Average Density of Root Hairs (AgDRH), Frequency of mycorrhization (F%), Intensity of mycorrhization (I%).

Altogether, our results suggest that root hairs development could play a weak role in root-adhering soil aggregation in pearl millet and that root architectural traits and AMF colonization rate have no significant impact.

### Genetic bases of rhizosheath formation in pearl millet

We previously reported the phenotyping of a panel of pearl millet inbred lines for root-adhering soil aggregation (Ndour *et al*., 2021). Briefly, a total of 1408 plants corresponding to 181 inbred lines were phenotyped and we recorded an almost four-fold variation in rhizosheath size (RAS/RT ratio), ranging from 7.4 (ICML-IS 11139) to 26.3 (ICML-IS 11084; Ndour *et al*., 2021). Here, we used these data to evaluate the heritability of root-adhering soil aggregation. A broad sense heritability of 0.72 was computed, suggesting that root-adhering soil aggregation is largely under genetic control. Altogether, these data indicate that root-adhering soil formation has high heritability and that a large genetic diversity exists in pearl millet.

We therefore analysed the genetic bases of root-adhering soil formation using association genetics. Out of the 181 inbred lines, 139 lines with good quality data for phenotype and genotype were retained to perform the association study. As a first step, we conducted a population structure analysis of the 139 lines (Supplementary Fig. S1) that confirmed the negligible genetic structure previously reported for this panel (Debieu *et al*., 2018). A total of 381,899 SNPs was used for the association analysis. We first calculated the least square means of the trait root-adhering soil aggregation (RAS/RT ratio) across the different experiments. The ratio ranged from 12.4 to 26.3 with an average of 18.0. The LFMM model for GWAS identified 53 significant SNPs (*p*-value < 0.0001) across the genome (Fig. 2A), defining 34 significant regions or QTLs considering windows of 50 kb up and downstream of significant positions to define significant regions. The proportion of phenotypic variance accounted for the most significant SNPs defining QTLs ranged from 9.2 to 15.6 % indicating that the corresponding QTLs had small phenotypic effect.

**Figure 2.**
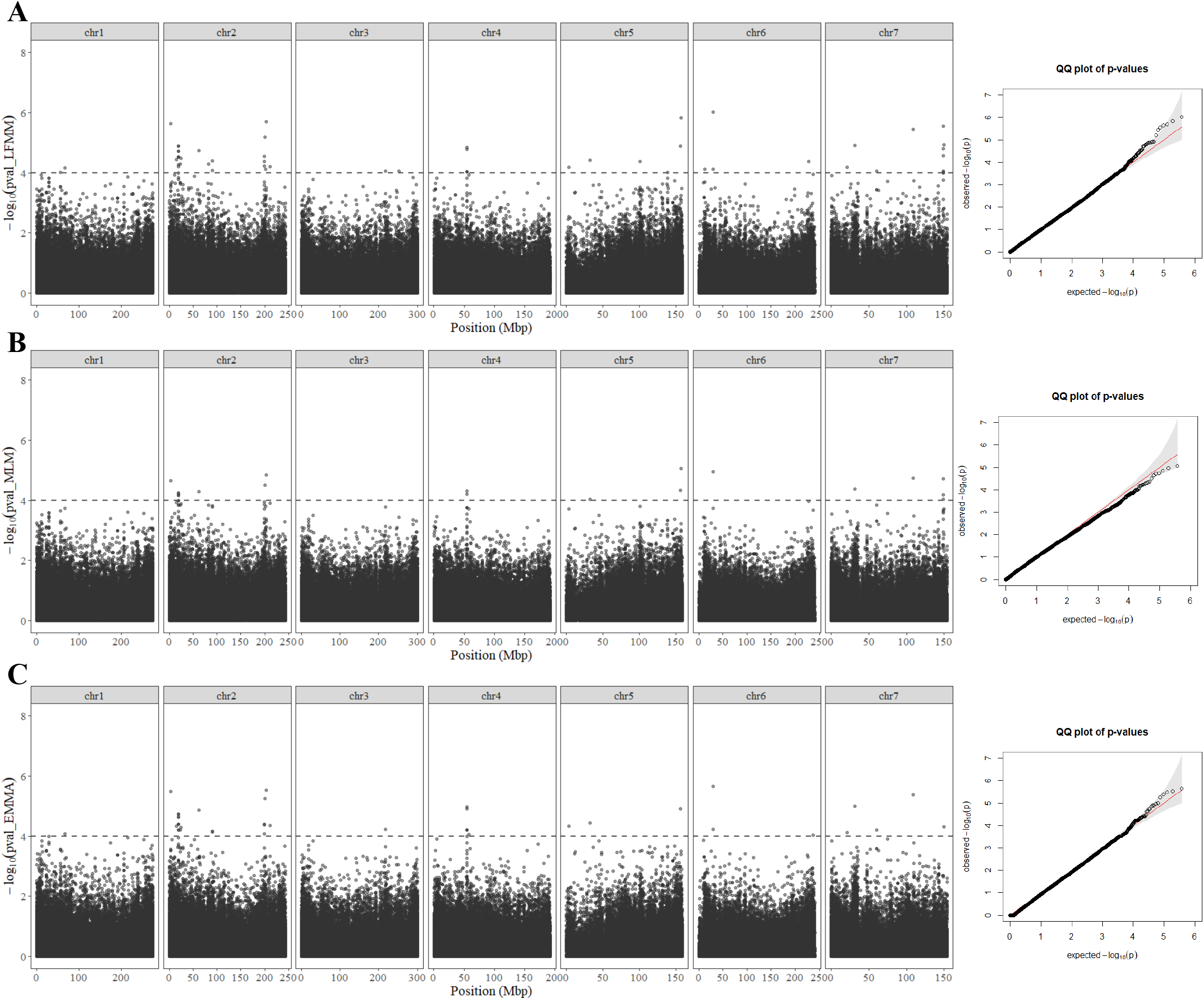
Genome-wide association studies (GWAS) for rhizosheath size in pearl millet. Manhattan plots and QQ plots obtained with three GWAS methods. **A)** Latent Factor Mixed Model or LFMM, **B)** Mixed linear model or MLM and **C)** Efficient Mixed Model Association or EMMA. Each Manhattan plot shows the –log10 *p*-value of the statistic (y axes) for each SNP position (x axes). The dashed line delimits the threshold for highly significant SNPs (*p*-value < 10-4).

We compared these results with two other GWAS methods (Fig. 2B&C). Thirty-nine of these SNPs included in 25 QTLs defined through LFMM were also found significant with EMMA, and 19 significant SNPs assigned to 14 QTLs using MLM model in GAPIT (Supplementary Table S2). Fifteen SNPs in 12 QTL regions were found significant across the three GWAS methods. Altogether, our GWAS analysis revealed 12 potential QTLs controlling root-adhering soil aggregation in pearl millet.

To back up our GWAS analysis, we performed bulk segregant analysis in a F2 population derived from a cross between two lines with contrasted RAS/RT phenotypes, ICML-IS 11139 (low RAS/RT) and ICML-IS 11084 (high RAS/RT; Fig 3A). F2 plants were phenotyped in five consecutive blocks together with the parental inbred lines. We confirmed the contrasted RAS/RT ratio of the parental lines with average values of 15.0 (ICML-IS 11139) and 32.7 (ICML-IS 11084, Fig. 3B). Ten individual F2 lines were dropped from the analysis leaving 547 F2 with RAS/RT ratio ranging from 1.6 to 54.8 and with an average value of 22.3. The phenotype distribution of the F2 was slightly skewed towards high values of RAS/RT ratio and showed a significant block effect (Supplementary Fig. S2). We used log transformation of RAS/RT ratio in our analysis of variance and selected lines with extreme values of the residual term for the bulks. The bulks consisted in two groups of 55 F2 lines each, with RAS/RT ratio average values of 11.0 for the low RAS/RT bulk and 38.2 for the high RAS/RT bulk (Supplementary Table S3). A total of 223.6 Mbp reads were mapped to the target enriched regions and used for SNP calling. We identified a group of 23,160 SNP variants (1.5 SNPs per 100 kb in average) between the bulks. The average sequencing depth was high with 887X and 863X in the small and high RAS/RT bulk respectively. The NGS-based BSA analysis revealed significant differences in the allele frequency of 380 SNPs at the 95% confidence interval (Table 2, Fig 3C). These SNPs defined five significant chromosome regions linked to the segregation of the RAS/RT ratio phenotype: three on chromosome 5 (RAS5.1, RAS5.2 and RAS5.3) and two on chromosome 6 (RAS6.1, RAS6.2; Table 2). The smallest genomic region defined corresponded to RAS5.3 with 10.6 Mbp and 41 significant SNPs. In contrast, the largest significant region corresponded to RAS5.2, with 45 Mbp and 307 significant SNPs.

**Figure 3.**
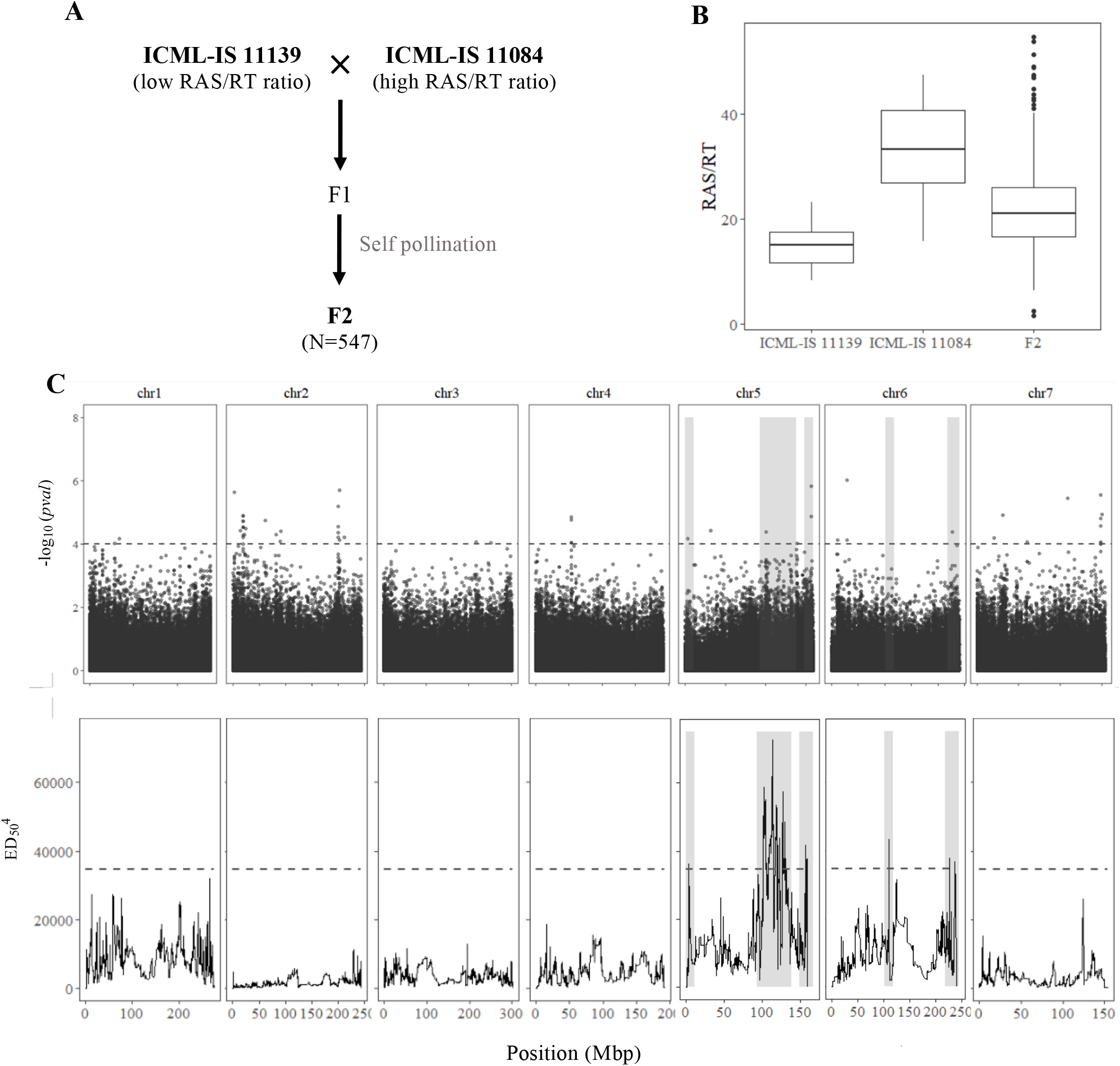
Genetic dissection of root soil aggregation in pearl millet by Bulk Segregant Analysis (BSA). **A)** Cross established for Bulk Segregant Analysis (BSA) between two pearl millet inbred lines with contrasted rhizosheath phenotype. **B)** Boxplot showing the distribution of RAS/RT ratio in line ICML-11139 (N=29), ICML-IS 11084 (N=27) and F2 population (N=547). **C)** Comparison between GWAS and BSA results. Top figure represents the Manhattan plot of the GWAS by LFMM ridge method (Caye *et al*., 2019). The x-axis corresponds to the position of the 381,899 SNPs identified by GBS in a group of 139 inbred lines. The vertical axes correspond to the –log10 *pvalue* of the statistic. The dashed line delimits the threshold for highly significant SNPs (*p value* < 10^-4^). Bottom figure shows the significant regions associated with root soil aggregation identified by BSA using bulks of contrasted F2 lines from a bi-parental cross. The plot shows the Euclidean Distance statistic profile (y axis) across the seven pearl millet chromosomes (x axis). The dashed line indicates the 95% confidence interval threshold for the localisation of significant regions. In both plots, the shaded area delimits the extent of the five significant regions identified by BSA and the overlap with significant SNPs identified by GWAS and the correspondence with the BSA peaks found.

**Table 2.** Significant genomic regions identified by Bulk Segregant Analysis (BSA) for rootadhering soil aggregation (i.e. RAS/RT) at the 95% confidence interval.

|  | Chr. | Peak position (Mbp)<sup>1</sup> | Region range (Mbp)<sup>2</sup> | Region length (Mbp) | Number sig SNPs |
| --- | --- | --- | --- | --- | --- |
| *RAS5.1* | 5 | 3.33 | 0 – 11.36 | 11.33 | 3 |
| *RAS5.2* | 5 | 112.88 | 92.95 – 137.95 | 45.01 | 307 |
| *RAS5.3* | 5 | 156.25 | 148.12 – 158.68 | 10.56 | 41 |
| *RAS6.1* | 6 | 110.41 | 102.08 – 118.41 | 16.33 | 16 |
| *RAS6.2* | 6 | 226.25 | 218.25 – 240.48 | 22.23 | 13 |

<sup>1</sup> Position of the most significant SNP in the region range
<sup>2</sup> Limits of the significant region considering the overlapping confidence interval of significant markers in the region

Interestingly, the range of four out of five BSA significant regions was found to overlay with the position of significant SNPs defined by GWAS (Figure 3C, Supplementary Table S2). Furthermore, the peak position of RAS5.1 on chromosome 5 is located 43 kb away from the SNP chr5_3282686 identified by GWAS (LFMM and EMMA). Likewise, the RAS5.3 spans through a genomic region containing two GWAS QTLs (LFMM, MLM and EMMA) located 113 kb and 244 kb away from the peak position of RAS5.3.

Altogether, the combination of GWAS and BSA analyses revealed genomic regions on chromosomes 5 and 6 controlling RAS/RT ratio in pearl millet.

### Comparison of gene expression in contrasted lines

To further analyse the genes involved in rhizosheath formation, we compared gene expression in ICML-IS 11139 (low RAS/RT) and ICML-IS 11155 (high RAS/RT) roots. Production and secretion of root exudates occur along the root system (Haichar *et al*., 2014), starting in the zone immediately behind the root tip (Schroth and Snyder, 1962). Similarly, root hair development occurs in the root tip. Thus, as these two processes seem to be the major determinants of root-soil aggregation in pearl millet, we hypothesized that genes controlling this trait might be preferentially expressed in the root tip. Phenotyping for RAS/RT ratio was performed at 28 days after planting, when the root system of pearl millet was made of one primary root and several crown roots possessing lateral roots (Passot *et al*., 2016). As crown roots make up most of the root system at this stage and to avoid noise due to sample heterogeneity (different root types), we therefore compared gene expression in the crown root tips (2 cm apex) of the two contrasted lines. RNAseq revealed 1270 genes with significant differences in gene expression between the two contrasted lines using three combined statistical tests (EdgeR, DESeq et DESeq2, *p*-value < 0.05; Supplementary Fig. S3). A gene ontology analysis on 742 genes with GO annotation out of the 1270 differentially expressed genes revealed a significant enrichment in GO terms associated with proteins involved in molecular interactions (GO:0043531, ADP binding with lowest *p*-value) and enzymatic reactions (GO:0016706, oxidoreductase activity for instance; Supplementary Table S4).

### Candidate genes analysis

We combined GWAS, BSA and gene expression analyses to identify candidate genes for RAS aggregation. We first focused our search for candidate genes in the QTL regions identified by GWAS that were coincident with regions of significance defined through BSA on chromosomes 5 and 6 (GWAS QTLs 5.1, 5.3, 5.5, 5.6 and 6.3). We assessed the annotated genes from the reference genome (Varshney *et al*., 2017) included in a 1 Mbp region centred around the most significant SNP position together with their expression data from the RNAseq experiment.

The most significant SNP marker in GWAS QTL 5.1 maps in chromosome 5 position 3,282,686 bp in an intergenic region between a cluster of four genes coding for glyoxylate reductase (*Pgl_GLEAN_10016760, Pgl_GLEAN_10016761, Pgl_GLEAN_10016762* and *Pgl_GLEAN_10016764*). Out of the four genes, one was differentially expressed in the contrasted lines for RAS/RT, the others showed a weak and variable expression level within the same genotype. Glyoxylate reductases are recycling enzymes that reduce glyoxylate to glycolate (Hoover *et al*., 2007). Interestingly, the glyoxylate cycle plays an important role in the synthesis of malate, which is a major metabolite excreted in root exudates (Fernie and Martinoia, 2009). This region also contains *Pgl_GLEAN_10016765*, a gene coding for an arginase with significantly higher expression in ICML-IS 11139 (low RAS/RT). Arginases metabolise arginine and provide nitrogen for the synthesis of other essential amino acids during plant development and stress response mechanisms (Siddappa and Marathe, 2020). Large variations in arginine concentrations have been associated with changes in root exudate composition in plants exposed to drought (Gargallo-Garriga *et al*., 2018).

The GWAS QTLs 5.5 and 5.6 are coincident with the same region of significance defined in BSA, RAS5.3. This 10.56 Mbp region contains 105 annotated genes in the reference genome. Interestingly, the most significant marker trait association for GWAS QTL 5.5 falls into a gene showing some homology to remorins (*Pgl_GLEAN_10037821*). Remorins are membrane proteins playing an important role in plant biotic interactions (Jarsch and Ott, 2011).

The most significant SNP in GWAS QTL 6.3 maps in chromosome 6 position 227,616,229 bp in a gene coding for a galactinol-sucrose galactosyltransferase that is more expressed in the low RAS/RT ratio line (*Pgl_GLEAN_10028942*). These are enzymes involved in the synthesis of raffinose (Lehle and Tanner, 1973), an oligosaccharide stored principally in seeds, roots and tubers. Accumulation of raffinose in wheat and tomato roots occurs in response to low P conditions (Sung *et al*., 2015; V.L. Nguyen *et al*., 2019). Raffinose also accumulates in roots of pea seedlings exposed to water stress (Lahuta *et al*., 2014). In addition, the secretion of this oligosaccharide in root exudates is linked to the complex biotic interactions in the rhizosphere (Fang and Leger, 2010; Liu *et al*., 2017).

We also looked for candidate genes associated with the most significant SNPs consistently identified by GWAS that do not coincide with regions of significance defined through BSA. The GWAS QTL 2.3 contains a cluster of five significant SNPs mapping in the same gene, *Pgl_GLEAN_10019483*, encoding an LRR receptor-like serine/threonine-protein kinase. This gene is strongly expressed in the root tip of both pearl millet lines. LRR receptor kinases are involved in the perception of signalling molecules (Chakraborty *et al*., 2019).

The GWAS QTL 6.2 consists of two SNPs markers on chromosome 6 mapping in an intergenic region between a cluster of four genes coding for acidic endochitinase (*Pgl_GLEAN_10020193,Pgl_GLEAN_10020194,Pgl_GLEAN_10020195* and *Pgl_GLEAN_10020196*), one of them with higher expression in the low aggregation line, the others with similar or weak expression in both lines. Endochitinase and chitinase-like proteins are defence related proteins with anti-fungal activity that are found in root exudates of different plant species (Nóbrega *et al*., 2005; Tesfaye *et al*., 2005; De-la-Peña *et al*., 2010).

In chromosome 7, we found a group of five significant SNP markers within a 74 kb range defining the GWAS QTL 7.5. These SNPs were close to a gene encoding a putative chloroplastic dicarboxylate transporter that exchanges malate for succinate, fumarate and 2-oxoglutarate (*Pgl_GLEAN_10006630*). All these are important components of root exudates. The region also contains a gene coding for an ABC transporter G family member (*Pgl_GLEAN_10006636*). ABC transporters are involved in the transport of root exudates (Badri *et al*., 2009; Baetz and Martinoia, 2014).

## Discussion

Here, we investigated root system architectural traits with potential impact on rhizosheath formation in pearl millet. The presence of root hairs is essential for rhizosheath formation but the impact of root hair length and density on rhizosheath size varies considerably between plant species (Brown *et al*., 2017). Studies in wheat and maize showed that root hair length is strongly correlated with rhizosheath weight (Delhaize *et al*., 2012; Adu *et al*., 2017). Similarly, in foxtail millet, increased rhizosheath formation was found related with the plastic response in root hair formation (increases in root hair elongation and density) in dry soils (Liu *et al*., 2019). This relationship was not as clear in crops such as barley (George *et al*., 2014). However, a recent study shows an increased rhizosheath formation in barley grown in drying soil associated with auxin-promoted growth of root and root hairs as a consequence of ABA accumulation (Zhang *et al*., 2021). In our study on pearl millet, we have identified a weak but significant correlation between rhizosheath formation, which is synonymous with root-adhering soil formation in this work, and root hair length (*p* = 0.005, *r^2^* = 0.077). This suggests that root hairs are involved in rhizosheath formation in pearl millet but that they play a limited role in our experimental conditions.

Root association with arbuscular mycorrhizal fungi (AMF) has been proposed to contribute to rhizosheath formation (Pang *et al*., 2017). In our study, we did not find any correlation between rhizosheath formation and AMF colonization rate in pearl millet suggesting that AMF colonization level is not an important driver in rhizosphere aggregation in this species.

Altogether, we hypothesise that rhizosheath formation or the aggregation of soil particles to the root in pearl millet is mainly driven by other traits. Root exudates and mucilaginous polymers released by root-associated microorganisms as well as the enzymatic activities linked to the crosstalk interactions occurring in the rhizosphere are prime candidates. Accordingly, the different orders of bacteria predominantly found in the rhizosphere of pearl millet lines with contrasted root soil aggregation suggests that the differences in rhizosheath formation could be linked to crosstalks between the plant and microbial community (Ndour *et al*., 2017, 2021). Further work will be needed to test this hypothesis.

In the current study, the large variation in rhizosheath size in a genetically diverse group of inbred lines revealed a high heritability value for the trait (H^2^=0.72). Although rhizosheath formation relies on a range of traits mainly related with root morphology and exudates, it has been found under genetic control in other cereal crops such as wheat (Delhaize *et. al*., 2015; James *et al*., 2016) and barley (George *et al*., 2014; Gong and McDonald, 2017), becoming a potentially interesting target trait for breeding (Ndour *et al*., 2020). Chromosome regions associated with rhizosheath size were identified in both crops, however few candidate genes underlying the QTL regions have been proposed. Interestingly, comparative evaluation of the multiple loci identified in these studies shows a lack of QTLs identified across diverse growing conditions suggesting, to some extent, a large QTL by environment interaction likely linked to the plasticity of rhizosheath formation.

Here, the combination of GWAS and BSA allowed the identification of four chromosome regions controlling rhizosheath size in pearl millet and ultimately some putative candidate genes based on gene annotations in the reference genome (Varshney *et al*., 2017). GWAS allowed the identification of 34 significant QTLs using the latent factor mixed model or LFMM (Caye *et al*., 2019) method. Many of these associations were confirmed using two other models for GWAS analysis (EMMA and MLM). The phenotypic variance explained (PVE) by these loci ranged from 11.2% to 14.7% suggesting that rhizosheath size as a complex trait determined by many QTLs of moderated effect in pearl millet. Consistently, studies in biparental and multiparental populations of wheat revealed several QTLs linked to rhizosheath formation with proportions of variation explained by QTLs around 5 to 10%. (Delhaize *et. al*., 2015; James *et al*., 2016). Nonetheless, one major QTL for the trait was also identified in wheat (James *et al*., 2016).

Few genetic studies have identified genes potentially involved in rhizosheath formation and their predicted functions were mainly linked to root system morphogenesis and growth. For example, root hair length is a major driver determining rhizosheath size in wheat and, accordingly, genes coding for basic helix-loop-helix family of transcription factors that are known to control root hair development were identified as potential candidates underlying a rhizosheath QTL in that species (Delhaize *et. al*., 2015). In barley, genes controlling cell division in root apical meristem at seedling stage and genes linked to tolerance to drought and cold were also identified as putative candidates underlying some genomic regions associated with rhizosheath size (George *et al*., 2014).

In contrast, in our study, candidate genes were mostly related to plant metabolism and transport. Combining BSA and GWAS analyses revealed five co-localizing QTL regions. Candidate genes in these QTLs regions were mostly linked with root metabolic activities such as the synthesis of compounds commonly found in root exudates. For instance, the glyoxylate reductase and the arginase identified as putative candidates for QTL 5.1 are involved in the reduction and storage of essential compounds (i.e., glyoxylate and nitrogen) required for metabolic processes that mediate the synthesis of organic acids like malate and the synthesis of amino acids, respectively (Igamberdiev and Eprintsev, 2016; Siddappa and Marathe, 2020). These are major primary metabolites of root exudates which variations in concentration can trigger plant adjustments to enhance root access and mobilisation of soil phosphate and nitrogen when these nutrients are limited (Carvalhais *et al*., 2011; Mora-Macías *et al*., 2017; Canarini *et al*., 2019). Further, these compounds have been found to promote chemotaxis of beneficial bacteria into the rhizosphere (Feng *et al*., 2018). In fact, a recent study showed how differences in malate concentration in root exudates impacted the composition of microbial communities associated with wheat and rice root systems (Kawasaki *et al*., 2021). Another potential candidate gene identified for QTL 6.3, a galactinol-sucrose galactosyltransferase, is involved in the synthesis of raffinose, an oligosaccharide which variations in concentration in root exudates has been found to favour root colonisation by rhizosphere microbes (Liu *et al*., 2017).

Our genetic analysis is therefore fully consistent with our analysis showing that root architectural, root hair and AM symbiosis traits are not or poorly correlated with rhizosheath formation in pearl millet, and with our expression study that shows that genes involved in plant metabolism are differentially regulated between lines with contrasted rhizosheath size. It is also consistent with previous research showing differences in the rhizosphere metabolic activity of pearl millet lines with contrasted rhizosheath size (Ndour *et al*., 2021). In this study, increased activity of enzymes such as chitinase and phosphomonoesterase was observed in the rhizosphere of pearl millet lines with larger rhizosheath (same contrasted RAS/RT lines used in the present work). We hypothesised that increased exudation in lines with larger rhizosheath size lead not only to an enhanced stability of root-adhering soil aggregates but also to a decrease of pH that could have stimulated these enzyme activities (Ndour *et al*., 2021). Moreover, the amount of root exudate and the function of these enzymes could also impact the rhizosphere microbial communities promoting rhizosheath formation and explain the difference found in microbiota diversity in contrasted pearl millet lines (Ndour *et al*., 2021).

In conclusion, our physiological and genetic analysis suggest a central role for root exudation (quantitatively or qualitatively) in the regulation of rhizosheath formation in pearl millet. Rhizosheath formation seems to be controlled by many QTLs with small effects. We identified several candidate genes controlling this trait and future work will focus on the validation and characterization of the molecular mechanisms regulating rhizosheath formation in pearl millet.

## Abbreviations

RAS: root-adhering soil

## Supplementary data

Figure S1. Ancestry estimation using the cross-entropy criterion.

Figure S2. Frequency distribution of the RAS/RT phenotype in the bi-parental population designed for BSA experiment.

Figure S3. Number of genes differentially expressed in two contrasted inbred lines.

Table S1. Spearman correlation between the different traits in 2018 and 2020 experiments.

Table S2. Significant marker-trait associations for root-adhering soil aggregation using 3 GWAS methods

Table S3. RAS/RT phenotype in the contrasted inbred lines and the bulks used for BSA.

Table S4. Top ten GO enriched terms in genes differentially expressed between contrasting inbred lines.

## Acknowledgements

This work was supported by the French National Research Institute for Sustainable Development (IRD), the French Agence Nationale de la Recherche (ANR grant RootAdapt ANR17-CE20-0022-01 to LL), the CGIAR Research Programme on Grain Legumes and Dryland Cereals (GLDC), the NewPearl grant in the frame of the CERES initiative by the Agropolis Fondation (AF 1301-015 to LL as part of the ‘Investissement d’avenir’ ANR-l0-LABX-0001-0l under the frame of I-SITE MUSE ANR-16-IDEX-0006), and by the Fondazione Cariplo (No FC 2013-0891).

## Authors contributions

YV, LC and LL conceptualized and supervised the research; LL acquired the funding. CFC, MND, PMSN, MD, AG, SP, CB, and MP carried out the measurements and performed formal analyses. All authors discussed and evaluated the data. CFC, MND, PMSN, AG, YV, LC and LL wrote the first draft of the manuscript; all authors revised the manuscript and gave final approval for publication.

## Data Availability

The data that support the findings of this study are openly available at the National Center for Biotechnology Information (NCBI). Genotyping (GBS) data are available in genbank under reference number PRJNA492967 (GWAS) and PRJNA769524 (BSA). RNAseq dara are available in the Gene Expression Omnibus (GEO) under reference GSE185425.

